# NASH patient liver derived organoids exhibit patient specific NASH phenotypes and drug responses

**DOI:** 10.1101/791467

**Authors:** S. McCarron, B. Bathon, D. Abbey, D. M. Conlon, D.J. Rader, K. Olthoff, A. Shaked, T.D. Raabe

## Abstract

To determine if patient liver derived organoids can in principle be a useful experimental model for non alcoholic staetohepatitis (NASH), we derived to our knowledge for the first time bipotent ductal organoids from endstage NASH patient livers, and for comparison from normal donor livers. We found that all tested NASH liver derived organoids exhibited a profound failure to dedifferentiate from the hepatic state back to the biliary state, consistent with the known poor regenerative capacity of NASH livers. Indeed, RNAseq on all tested NASH organoid populations confirmed down regulation of multiple cell cycle pathways. NASH liver derived hepatically differentiated organoids can slowly expand as monolayers, significantly simplifying microscopic quantitation: The monolayers show variable, but overall significantly increased lipid droplet accumulation in response to free fatty acids. Transcriptome analysis of NASH organoids reveals strong upregulation of a wide variety of pro inflammatory pathways in a NASH patient specific manner. Surprisingly, NASH liver derived organoids are highly diverse not only regarding their cytochrome cytochrome p450 metabolism and inflammatory response, but also react differentially to known antisteatotic, anti inflammatory and antifibrotic drugs, raising the possibility of using NASH patient liver biopsy derived organoids for personalized drug screening and therapy.

**One Sentence Summary:** NASH patient liver derived organoids replicate NASH liver phenotypes in a patient specific manner and exhibit profound differences in their response to drugs that are currently used in NASH clinical trials.

## Introduction

Nonalcoholic fatty liver disease (NAFLD) is the most prevalent chronic liver disease world wide and is closely associated with type 2 diabetes mellitus (T2DM) (*1*, *2*). While NAFLD, which is characterized by excessive liver fat accumulation, usually defined as fat exceeding 5% of the liver weight, can be benign over many years and is indeed somewhat reversible though control of diet and through exercise, a significant subset of NAFLD patients develop nonalcoholic steatohepatitis (NASH), a late stage form of NAFLD characterized by inflammation that often progresses to cirrhosis and hepatocellular cancer (HCC). Indeed, NASH is rapidly becoming the leading cause for end-stage liver disease or liver transplantation (*3*). There is considerable urgency regarding the development of effective anti NASH drugs, as there are currently none available except in clinical trials, and as the prevalence of diagnosed NASH could reach 18 million by 2027 in the US, Japan, and the EU 5 (England, France, Germany, Italy, and Spain) (*4*). While dietary caloric restriction and exercise can be somewhat effective in a subset of patients they clearly do not work for most patients, underscoring the dire need for pharmacotherapy.

At present it is difficult to predict which steatosis patient will go on to present severe inflammation, fibrosis and cirrhosis, although such information would be very valuable for timely treatment and prevention of these severe conditions. In general it is assumed that more than one type of stimulus is needed to cause severe liver injury. For example a high fat diet combined with a genetic predisposition, or a specific probiotic gut environment combined with high caloric diet could lead ultimately to activation hepatic stellate cells which then produce excessive amounts of extracellular matrix causing fibrosis, further inflammation and eventually irreversible and terminal cirrhosis, which can only be treated by liver transplantation.

While animal models of liver disease, including NASH, have been invaluable for definition of important liver disease mechanisms, it is becoming increasingly clear that many aspects of liver disease are unique to primates, including humans, and thus should be studied using human tissue. One example is the recent failure of a DGAT2 inhibitor to lower plasma triglyceride levels in rhesus monkeys while it was very effective in mice (*5*).

Recently, mouse and human induced pluripotent stem cell (iPSC) or mouse fibroblast derived hepatocyte like cells (iHeps), have been described and are used by many groups for liver disease modeling including after transplantation into rodents, although they do require several derivation and differentiation steps (*6*–*9*). In addition, derivation of non-integration and feeder free iPSC themselves from human blood takes routinely 3-6 months.

The DNA methylation pattern in NASH patient livers have recently been shown to be strikingly distinct from those of normal livers (*10*, *11*) and a large part of these epigenetic changes could be due to highly liver specific environmental factors such as the type of nutrients, xenobiotics and micoorganisms that enter the liver at very high concentrations through the portal tract where they are subjected to major metabolic modification. <u>iPSC derived iHeps are thus less likely to reflect the methylome of the liver compared to hepatically differentiated stem cells that were directly derived from the liver. In addition, the inherent genetic instability during iPSC derivation could introduce unforeseen genetic changes (*12*). Together, these factors may compromise interpretation of functional data from iPSC derived iHeps, especially regarding phenotypes that are controlled by the methylome</u>.

In 2015, the Clevers group presented a method for isolation of proliferating bipotent ductal organoids directly from normal adult human donor liver, mostly acquired from small biopsies of donor livers that were otherwise used for transplantation. These organoids are genetically an order of magnitude more stable than iPSC (*12*), can be cultured over a period exceeding one year (*12*, *13*) and can be cloned and genetically manipulated (*13*). Upon simply changing the medium from expansion medium to differentiation medium they stop dividing and acquire many properties of adult human hepatocytes. They are also capable of engraftment in immune compatible fah−/− mouse liver, after splenic injection (*12*) and their epigenome should closely reflect the epigenome of the liver they were derived from.

However, it is not known whether end stage NASH liver explants can also give rise to such bipotent ductal organoids, and if so, whether such organoids would at least in part recapitulate the underlying disease of the liver they were derived from. We present here data showing that severely injured NASH liver can still give rise to long term expandable bipotent ductal organoids which - after hepatic differentiation - can functionally recapitulate important aspects of the pathology of the liver from which they were derived.

## Results

### Bipotent ductal organoids can be derived from biopsy-size amounts of NASH patient liver or normal human liver

For ductal organoid isolation we chose to use a recently published protocol designed for human donor liver as a starting point. Using several modifications (see method section), we were able to derive bipotent ductal organoids from end stage patient liver explants that were otherwise discarded as a byproduct of liver transplantation. To our surprise, so far we have never failed to derive long term expanding organoids from any of the explant NASH liver samples we have tried (a total of 8 as of July 2019; see also supplemental Figure 1), although the initial yield of organoids varied from explant to explant, as it does for organoids from donor liver. While we initially used 300mg - 500mg of explant tissue for organoid derivation, we later showed that we can readily use ~100mg explant liver as starting material, with an acceptable reduction of yield compared to 300-500 mg. Importantly, in 3 out of 3 such tries, we obtained organoids of identical morphology and growth properties from both the 300-500mg and the ~100mg samples done in parallel. We note that 100mg is a sample size that can be provided by percutaneous liver biopsy. For comparison we also isolated organoids from normal human liver. Indeed, in all cases we were able to derive viable organoids from 50-100mg of donor liver tissue. Liver tissue from normal living liver donors (2 total) produced higher densities of organoids compared to donor livers (7 total) that were obtained through the Organ Procurement and Transplantation Network (OPTN). For OPTN livers organoid yield varied, likely at least in part due to the variation in quality of the transported livers. For the purpose of this paper we compared 6 NASH liver derived organoids to 3 donor liver derived organoids, two of which we obtained from living donors (see Table 1 and supplemental Figure 1).

**Table 1.**
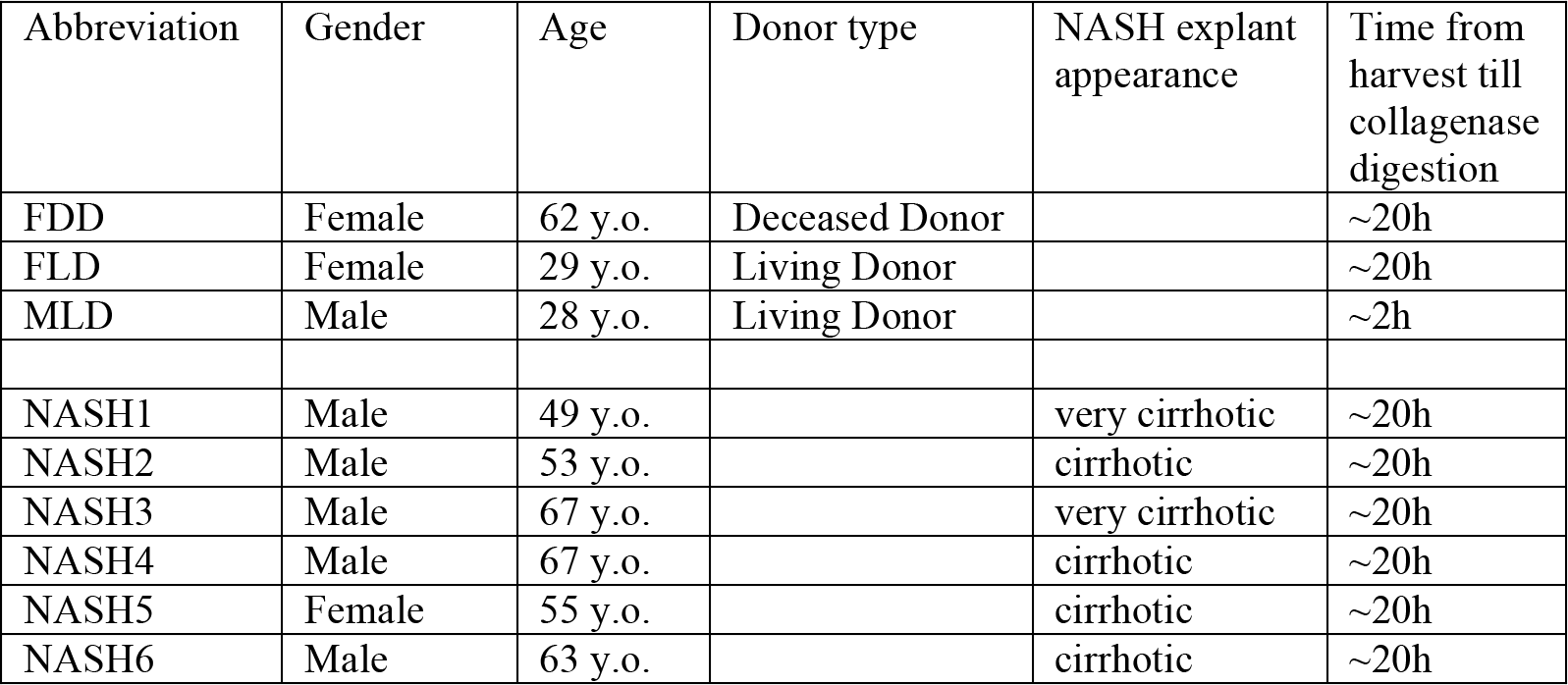
NASH liver explants and normal donor liver biopsies used in this study.

| Abbreviation | Gender | Age | Donor type | NASH explant appearance | Time from harvest till collagenase digestion |
| --- | --- | --- | --- | --- | --- |
| FDD | Female | 62 y.o. | Deceased Donor |  | ~20h |
| FLD | Female | 29 y.o. | Living Donor |  | ~20h |
| MLD | Male | 28 y.o. | Living Donor |  | ~2h |
| NASH1 | Male | 49 y.o. |  | very cirrhotic | ~20h |
| NASH2 | Male | 53 y.o. |  | cirrhotic | ~20h |
| NASH3 | Male | 67 y.o. |  | very cirrhotic | ~20h |
| NASH4 | Male | 67 y.o. |  | cirrhotic | ~20h |
| NASH5 | Female | 55 y.o. |  | cirrhotic | ~20h |
| NASH6 | Male | 63 y.o. |  | cirrhotic | ~20h |

### Ductal organoids isolated human NASH explant liver exhibit developmental delay and characteristic growth patterns

Compared to normal liver derived organoids NASH patient livers exhibit varying degrees of developmental delay after isolation, and all exhibit longer division times even after many passages (see supplemental Figure 2 for an example). Indeed, all organoid lines from normal liver as well as explant NASH livers continued to grow without significant change in growth pattern at all times of culture even at the highest passage numbers (for details see supplemental Figure 1 and Table 1). This could not be explained by normal variation due to trivial factors such as amount of starting material, variations in organoid isolation procedure, or variation in media preparations. The most significant delay of organoid growth was seen for NASH patients 2 and 4 (see supplemental Figure 1C). Significant differences can also be seen in the overall shape as well as the pattern of cells of hepatically differentiated organoids. For example patient 2 consistently generates irregularly shaped organoids compared to normal liver derived organoids (see supplemental Figure 1).

### Organoids from adult normal, but not human NASH explant liver, can repeatedly and efficiently cycle between the biliary and the hepatic state

Since previous work using sophisticated lineage tracing in mice indicates that hepatocytes can dedifferentiate towards a proliferative state *in vivo* and convert to cholangiocytes in the presence of genetically ablated cholangiocytes (*14*) and choangiocytes can convert to hepatocytes in the presence of chemically injured hepatocytes (*15*) we wondered if addition of expansion medium to differentiated cells just after splitting would allow the cells to re enter the proliferative state.

Surprisingly, hepatic organoids from normal human donors cultured for more than two to three weeks in differentiation medium were able to very efficiently exit the hepatic state after being split into fresh BME matrix and expansion medium. Remarkably, the degree of dedifferentiation was such that growth rate and morphology comparable to those of ductal organoids grown exclusively in expansion medium were observed as early as one week after the split. Next, we wondered if organoids from liver explants would behave similarly. To our surprise, all of the explant derived organoids we tested so far showed a very significant reduction of their capability to regenerate from the hepatic state back to the biliary state (Figure 1B). We note that irrespective of the cause of the end stage liver disease, all tested organoids from explant liver are compromised in their ability to dedifferentiate quickly, although at varying degrees (Figure 1B).

**Figure 1.**
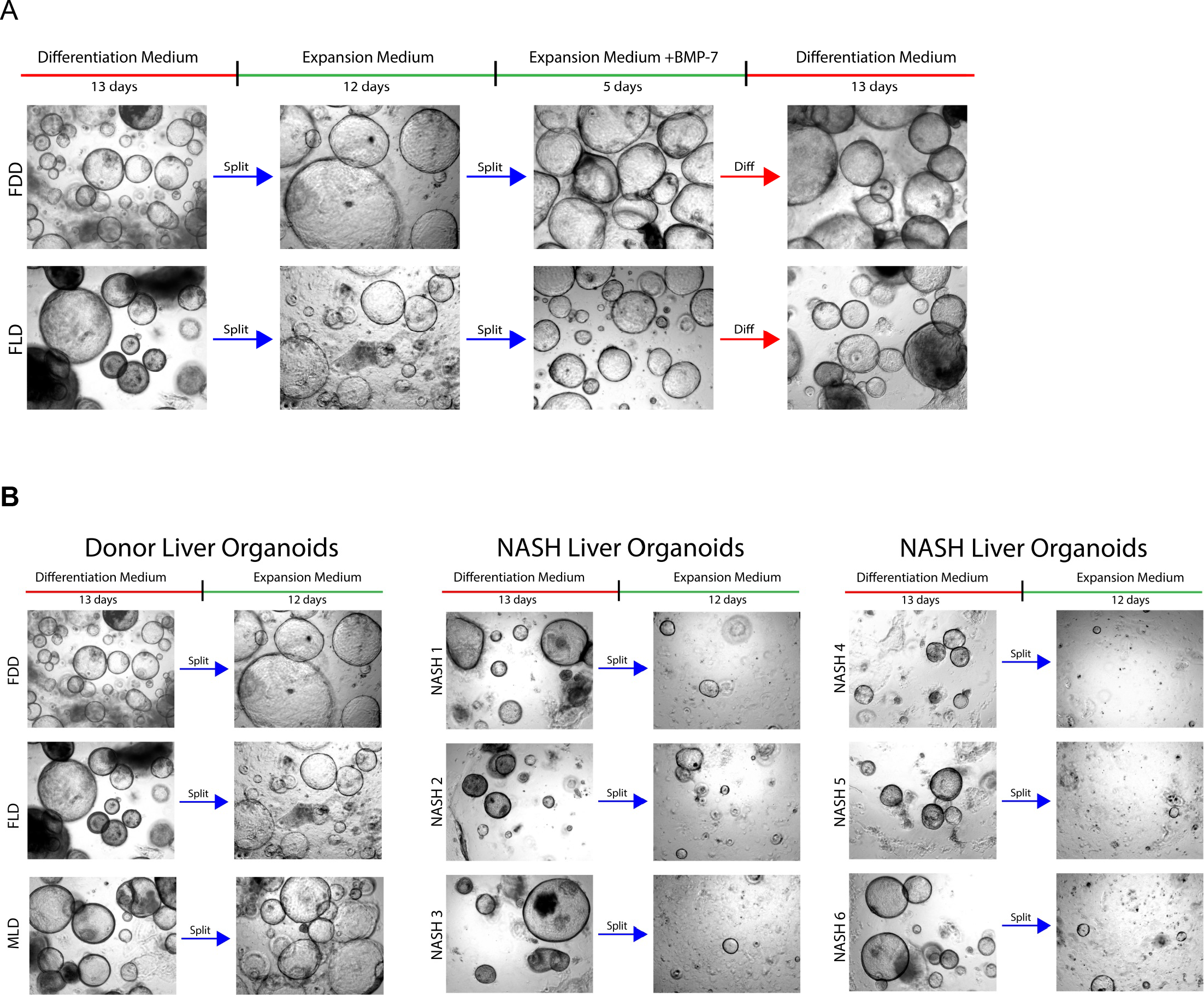
Organoids from adult normal, but not human NASH explant liver, can repeatedly and efficiently cycle between the biliary and the hepatic state. **A.** All organoids from normal liver repeatedly cycle between hepatic and biliary states. Expanding female donor organoids were differentiated to the hepatic state for 13 days. Organoids were then split into expansion medium to dedifferentiate back to the biliary state. After subsequent expansion and passaging the same cells were redifferentiated to the hepatic state a second time. Redifferentiation required the same amount of time as the first differentiation. **B.** Failure of all hepatically differentiated organoids form NASH liver to dedifferentiate back to the biliary state. Normal and NASH liver organoids were differentiated in parallel to the hepatic state for 13 days. Organoids were then split and seeded into expansion medium. All organoids from NASH liver exhibit a block of dedifferentiation to the biliary state.

### Hepatically differentiated organoids isolated from human NASH explant liver exhibit decreased LDL uptake and increased oleic acid induced lipid accumulation

Unexpectedly, we noticed for all 3 of our donor organoids and all 6 NASH organoids that during differentiation cellular monolayers would slowly extend beyond the BME matrix drop at various average distances, reaching the wall of the well in some cases, depending on the organoid population. Therefore not all organoid cells are strictly non proliferative during differentiation - instead they grow as flat cells and multiply slowly during differentiation (see Figure 2A). Indeed there is clear slow growth of flat cells for at least 4 weeks after differentiation. Since we wanted to analyze various metabolic functions in differentiated organoid we reasoned that the flat cells may provide an excellent opportunity to quantitatively analyze differentiated cells by live fluorescent microscopy.

**Figure 2.**
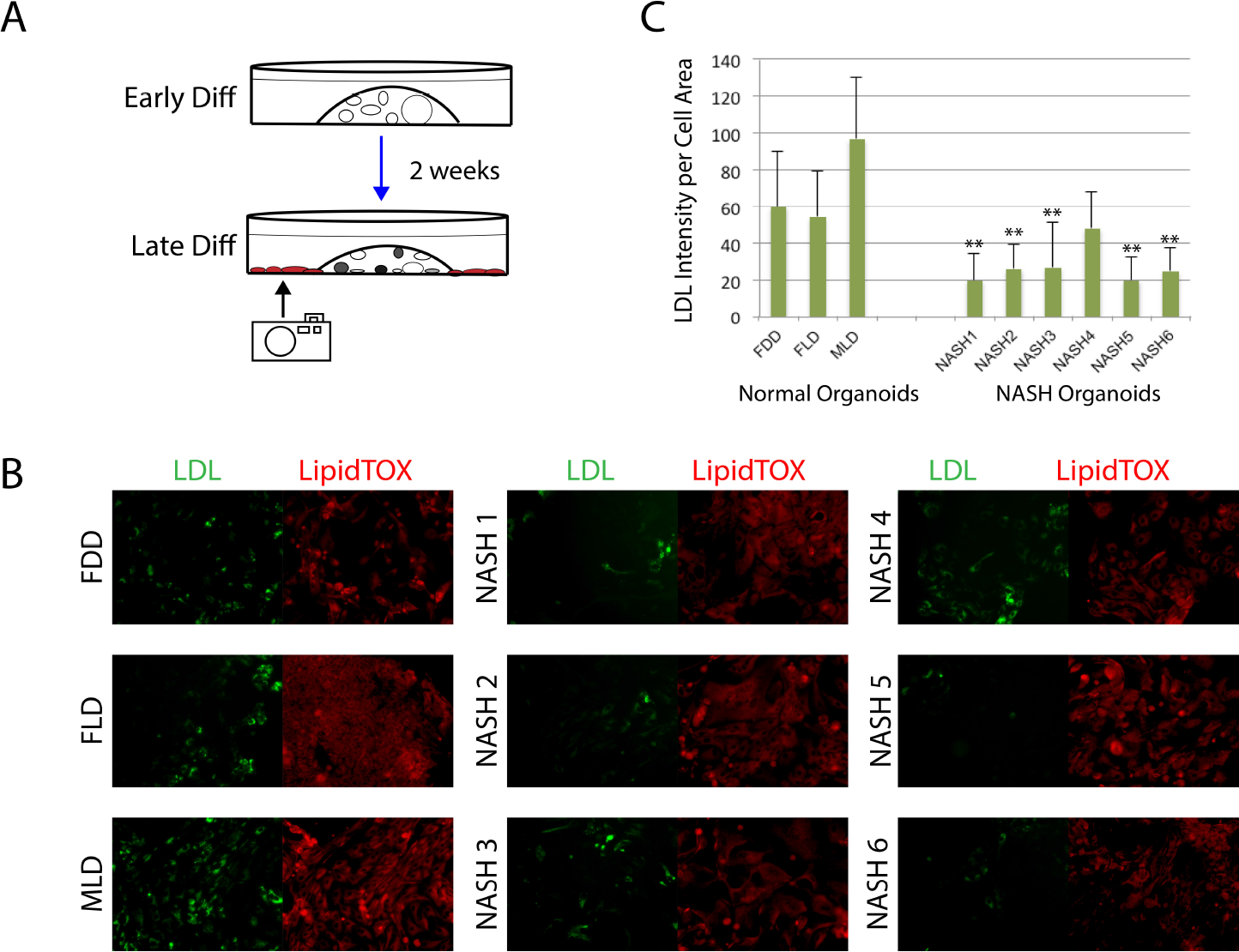
Hepatically differentiated organoids isolated from human NASH liver explant exhibit lower LDL uptake. **A.** Schematic of increase in flat cell outgrowth throughout differentiation. **B.** NASH organoids exhibit lower LDL uptake compared to donor organoids. Normal donor and NASH organoid flat outgrowth cells were double labeled with green fluorescent LDL and red fluorescent LipidTOX on day 13 of differentiation. After overnight incubation, flat cells were imaged (10 images for each organoid flat cell population). One representative image is shown for each cell population: LDL= green (left) and LipidTOX = red (right) **C.** Quantification of LDL uptake per cell area was done using NIS elements software (n=10 for each data point). Cell area was measured by over exposing lipidtox stain which reveals 100% of cell areas, even those with low lipid levels. P values are for each NASH patient vs the same normal donor (FDD): ** p<0.006.

For these monolayers, we devised a live fluorescent microscopy double labeling assay measuring simultaneously lipids as well as uptake of fluorescent LDL and show that LDL uptake is downregulated in organoids from NASH livers (see Figure 2). Visual inspection indicated consistently that flat cells as well as 3 D organoids appeared to be similarly stained for LDL uptake and lipid accumulation. Further, we devised a triple labeling assay to measure lipid droplet accumulation in response to oleic acid, number of nuclei and apoptosis (see Figure 3 and supplemental Figure 3). To determine lipid intensity per nucleus we counted the nuclei in each microscope image taken. NASH organoid flat outgrowth cells accumulate significantly more lipids than normal organoid flat outgrowth cells in response to 2mM oleic acid over a 24h period. Importantly, there are highly significant variations among individual NASH organoid flat outgrowth cells.

**Figure 3.**
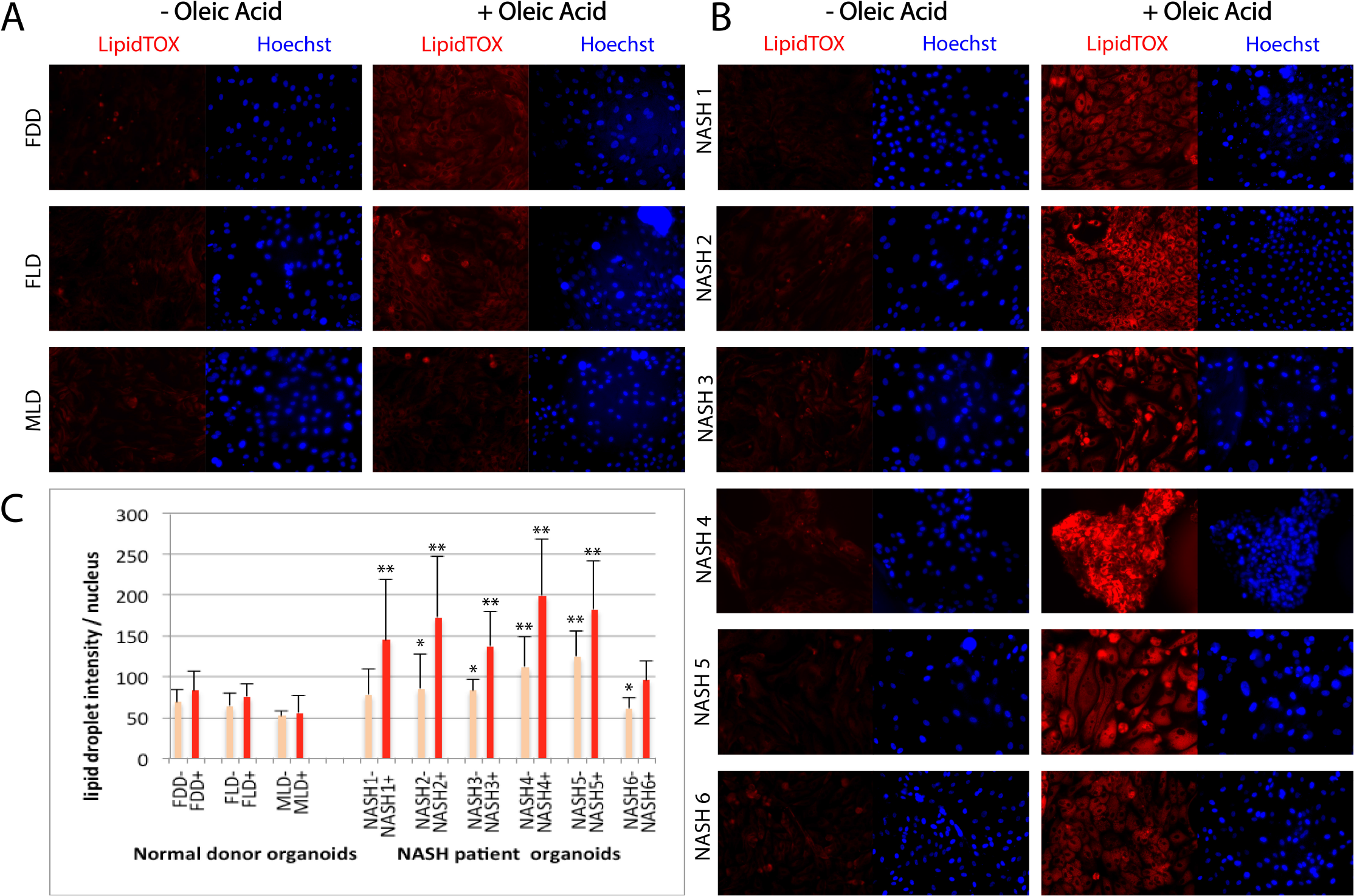
Hepatically differentiated NASH organoids exhibit increased free fatty acid induced lipid accumulation. **A and B.** After 2 weeks of differentiation, organoids either received no oleic acid or 2mM oleic acid for 24 h. Organoid flat outgrowth cells were double labeled with red fluorescent LipidTOX (lipid accumulation) and Hoechst dye (nuclei). Representative images of each organoid population (from a total of 10 images) from the same 3 normal donors (A) and 6 NASH patients (B) that were also tested for LDL uptake (see Figure 2) are shown. **C.** NIS imaging software analysis of lipid amount per nucleus (n=10 for each data point) shows that most NASH flat cells exhibit higher lipid uptake compared to donor flat cells. Orange bars: no oleic acid. Red bars: 2mM oleic acid. P values are for all NASH patients vs the same normal donor (FDD): *p<0.03, ** p<0.005.

In contrast, for apoptosis we noticed much less heterogeneity between individual NASH patients. We also did see a distinct difference between 3D organoids and flat cells for both normal organids and NASH organoids: Characteristic and strongly apoptotic cell clusters were found only among 3D organoids but not among flat cells (see supplemental Figure 3). However, these highly discrete ‘foci’ of a few to > 100 cells within the 3D organoids did not account for more than about 10% of the total 3D organoid area (see supplemental Figure 3).

### Screening for drugs that functionally alleviate the regenerative block and steatosis in organoids from NASH patients

Since lipid droplet accumulation is known to be an important initial trigger for the inflammatory response and hepatic stellate cell activation that causes fibrosis, and because there are several anti steatotic, anti inflammatory and anti fibrotic drugs in current clinical trials we reasoned that the functional heterogeneity in lipid metabolism (see Figure 3) may also be reflected in differential response to such drugs and may account for the variability of responses often found in clinical trials.

Numerous detailed studies have proven that the Hippo pathway is an essential regulator of liver regeneration and that the transcriptional co factor YAP, its main effector, is strictly required for liver regeneration after liver injury, such as partial hepatectomy or chemical injury (*16*–*19*) and YAP agonists induce proliferation of liver cells. We therefore tested the capability of known YAP upregulating regulating drugs as well as anti-inflammatory / anti fibrotic drugs currently in NASH clinical trials, for their ability to induce growth in monolayers of the hepatically differentiated NASH patient derived organoids table (see Figure 4). We chose drugs known to activate YAP in hepatocytes, namely XMU-MP1 (*20*) and LPA (Lysophosphatidic acid) (*21*). XMU-MP1 has also been shown to alleviate steatosis and fibrosis in mice (*20*). In addition we tested the thyroid receptor agonists T3 (Triiodothyronine) and its derivative MGL3196, the latter in promising NASH clinical trials (*22*). Further, we tested the anti inflammatory / anti fibrotic drugs Elafibranor (agonist of the liver specific transcription factor PPARα/δ and Obeticholic Acid (agonist of the bile acid receptor FXR) which are currently in clinical NASH trials (*23*–*26*). For further comparison we used simvastatin (a statin) and the potent direct YAP inhibitor verteporfin. Our data indicate that XMU-MP1 shows consistent activation of cell growth and verteporfin consistent inhibition, while all other drugs show profound differences in the proliferative response among the NASH patients (see Figure 4 for detailed results).

**Figure 4.**
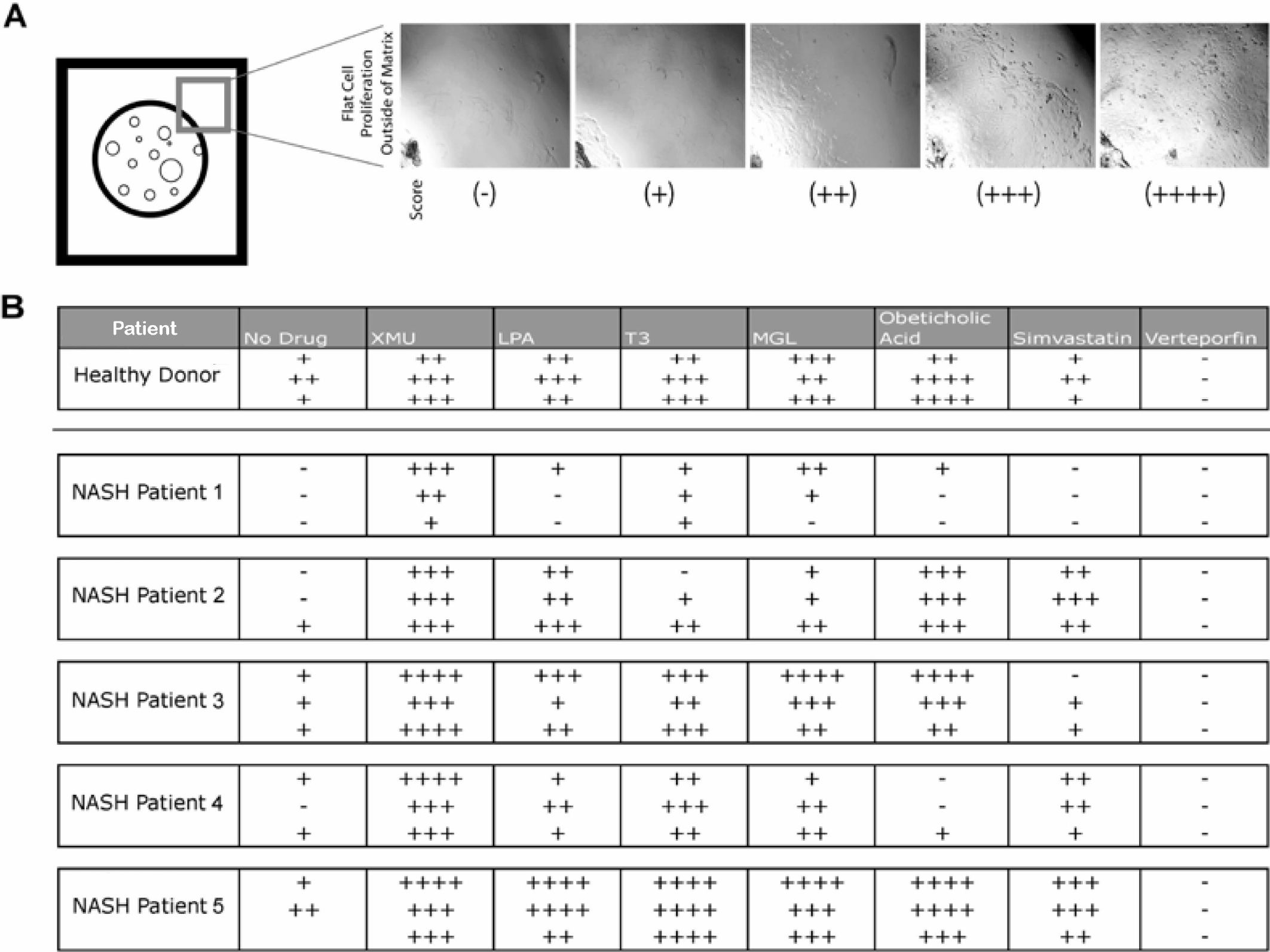
NASH liver derived organoids exhibit a wide spectrum of drug responses to clinically relevant drugs. **A.** Establishment of a visual scoring system for flat cell outgrowth beyond the 3D BME matrix area **B.** The YAP activating drugs XMU-MP1, LPA (Lysophosphatidic acid) and T3 (Tri-Iodothyronine) and its derivative MGL3691, as well as the anti inflammatory / anti fibrotic drugs, Elafibranor and Obeticholic Acid (the last 2 are currently in clinical NASH trials) and for comparison simvastatin (a statin) and the potent YAP inhibitor verteporfin were added to 2 week long differentiated organoid populations from donor FDD as well as NASH patients 1-5 (table1) and incubated an additional 2 weeks in differentiation medium. Experiments were done in triplicate in 48 well plates. The extent of flat cell outgrowth was measured by visual inspection of the wells as shown in A.

For high through put drug screening (HTS) it is important to use 96 well plates, or better 384 well plates. Towards this purpose, we devised a way to culture in 96 well plates exclusively flat monolayers derived from NASH organoids that grow under differentiation conditions (see Figure 5). Instead of differentiating already expanding 3D organoids by changing from expansion to differentiation medium, we put expanding organoids directly after mechanical disruption into differentiation medium containing 5% BME matrix which is soluble under these conditions. Indeed, cells readily attached to the bottom and subsequently formed slowly but continuously growing monolayers without going first through the 3D stage.

**Figure 5.**
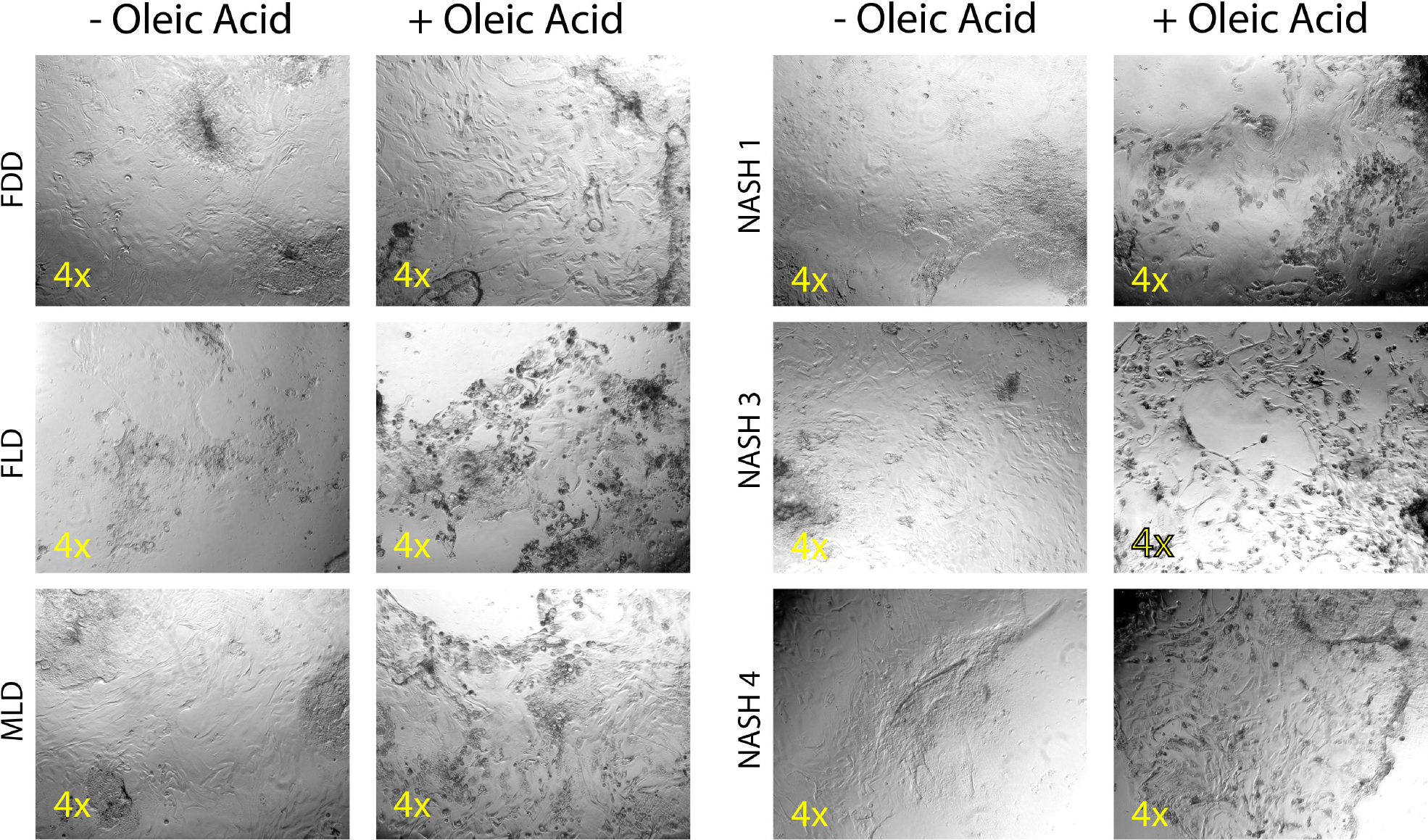
NASH liver derived hepatically differentiated organoids can be adapted to exclusive growth as monolayer in 96 well plates. The monolayer 96 well platform confirms the oleic acid induced lipid accumulation done in the 48 well 3D cultures from Figure 3. 4x phase contrast images show differentiated flat cells for donors FDD, FLD and MLD as well as NASH patients 1,3, and 4 with or without 2mM Oleic Acid addition for 24h.

### Human NASH liver explant derived organoids exhibit highly distinct transcriptomes

Prompted by the clear morphological and functional distinctions of NASH explant vs normal donor liver organoids we described above in detail, we decided to probe the transcriptome of hepatically differentiated organoids from the 3 normal donor livers and the 6 NASH explant livers in Table 1. They were all grown for 5 or 6 passages in parallel and then each split into 3 identical wells of a 24 well plate and differentiated for 13 days by simply changing from expansion medium to differentiation medium as described in the methods and supplemental Figure1. We performed RNA seq as described in the methods for 3 × 9 wells, resulting in 27 libraries. As shown in figure 6 A, library qualities are good as all triplicate libraries cluster together. Volcanoplots show highly distinct and very pronounced transciptome changes for all 6 NASH patients (Figure 6B). Combined transcriptomes of organoids from NASH patients 1-6 vs. combined of transcriptomes of organoids from normal donors FDD, FLD and MLD show strong overall upregulation of several metabolic pathways including cytochrome p450 pathways, and significant downregulation of many cell cycle – related pathways. While the former is surprising (see discussion), the latter is consistent with the poor regenerative ability of the NASH organoids (Figure 1).

**Figure 6.**
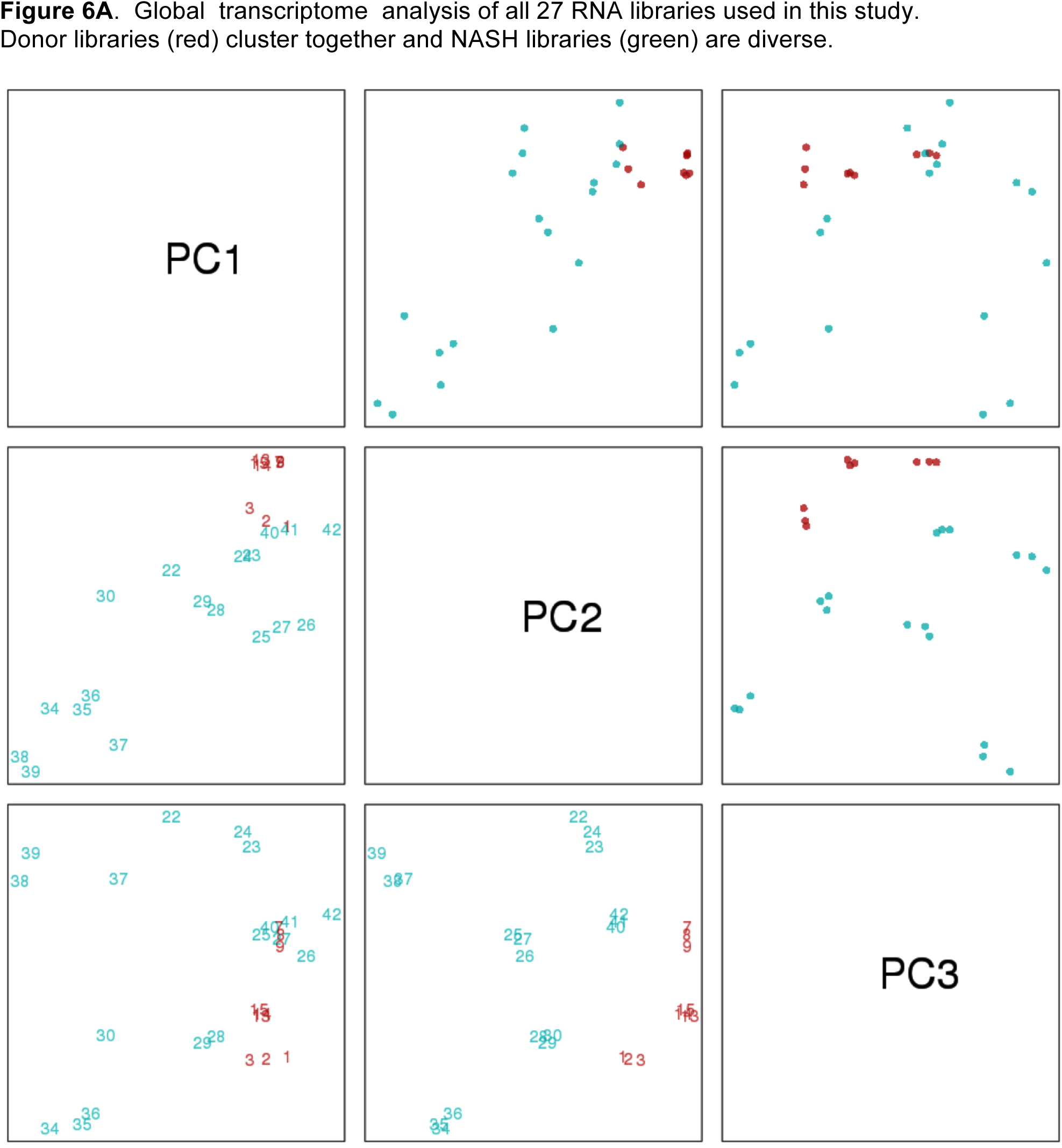
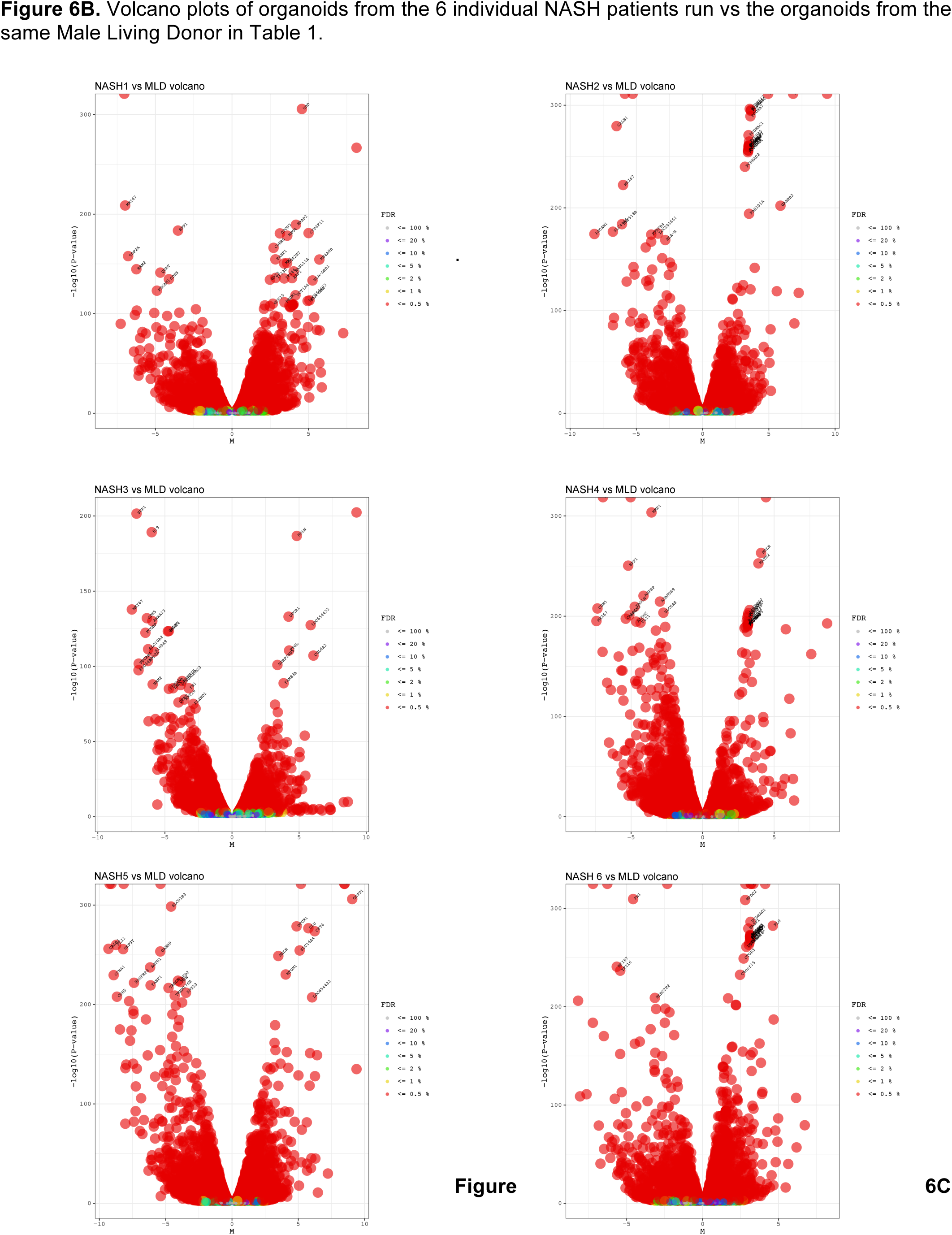
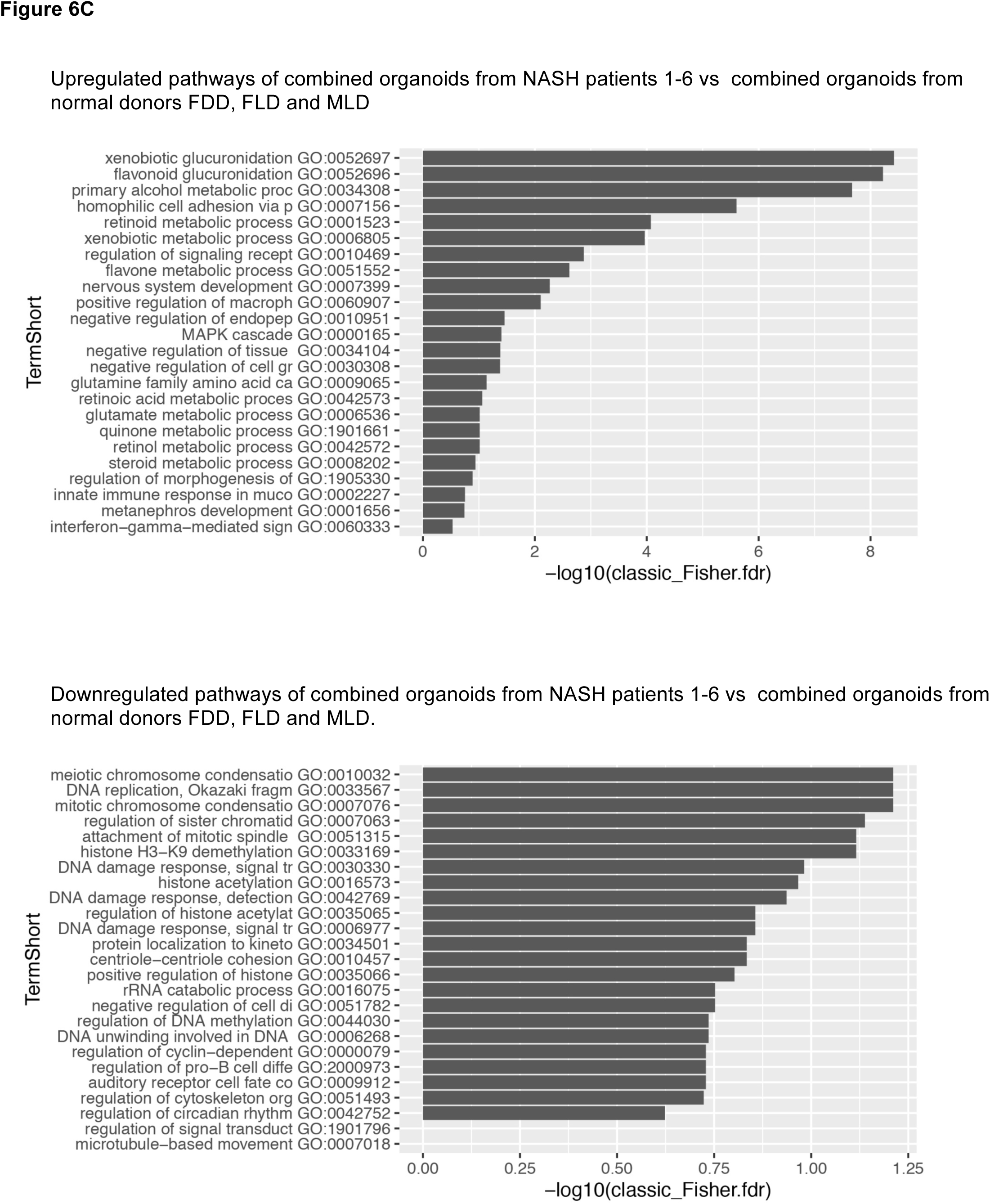
Hepatically differentiated organoids isolated from human NASH liver exhibit distinct transcriptomes. **A.** Global transcriptome analysis (using 6 different algorithms) of hepatically differentiated organoids from each of the 3 donors in Table 1, each grown in triplicate (red numbers show all 9 donor libraries) and hepatically differentiated organoids from each of the 6 NASH patients from Table 1 grown in triplicate (green numbers show all 18 NASH libraries). Donor organoids cluster together while NASH organoids are much more diverse from each other. **B.** Volcano plots of organoids from the 6 individual NASH patients run vs the organoids from the same Male Living Donor in Table 1. **C.** Significantly over expressed (upper panel) and under expressed pathways (lower panel) in the combined organoid transcriptomes of organoids from all 6 NASH patients vs the combined transcriptomes of all normal donor organoids from Table 1.

We next determined the upregulated pathways among individual NASH organoid transcriptomes vs the same Male Living Donor organoid transcriptomes (see Figure 7). To our surprise NASH organoids 1,2 and 5 show extremely strong upregulation of multiple Xenobitic and other metabolic pathways, while NASH oganoids 3,4 and 6 show little, if any, similar upregulation. NASH organoids 1,2,3 and 4 and 6, but not 5, show a broad variety of strongly upregulated proinflammatory pathways. Lipid pathways are over all not strongly regulated, indeed, the transcriptomes were assessed in the absence of stimulatory free fatty acids.

**Figure 7.**
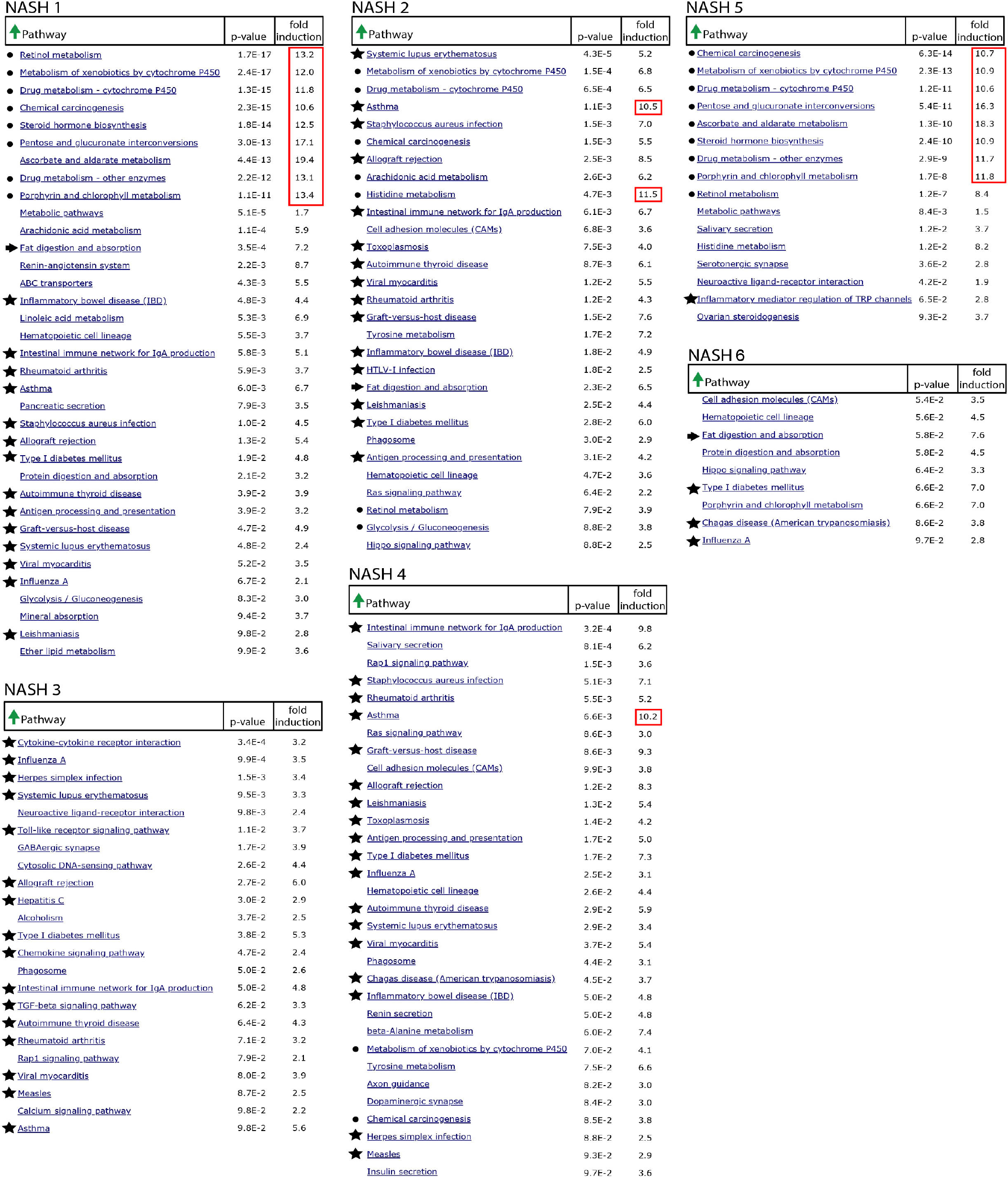
Over expressed pathways vary strongly among the 6 individual NASH patient derived hepatic organoids vs the same Male Living Donor hepatic organoids (see Table 1 and suppl. Figure 1). RNA seq data from the libraries in Figure 6A were entered into the online David 6.8 Analysis Wizard (NIAID/NIH) to determine over expressed pathways in NASH organoids ranked by p-values. Black dots denote pathways of xenobiotic, cytochrome p450, retinol and steroid metabolisms; stars mark proinflammatory pathways and arrows show lipid pathways. More than 10 fold over expressed pathways are encircled by a red box.

## Discussion

We derived to our knowledge for the first time bipotent ductal organoids directly from NASH patient liver explants. Indeed only 100mg of liver tissue was needed for derivation of organoids from both, NASH patient liver and Donor liver. 100mg of liver tissue is comparable with the amounts that can be obtained by routine needle biopsy. This raises the possibility of routine liver biopsies for derivation of organoids. Indeed, routine liver biopsy remains the gold standard for diagnosis of NASH and if performed optimally poses minimal risk to the patient (*23*, *27*, *28*). Indeed needle biopsies have recently been reported to be sufficient for derivation of liver cancer organoids from patient liver (*29*) despite the fact that they actually grow more slowly in organoid culture than normal liver organoids (*30*).

We noticed that normal donor liver derived organoids are not only bipotent but can <u>repeatedly</u> convert from the biliary state to the hepatic state and back to the biliary state, closely reflecting the situation *in vivo*, as several careful lineage tracing studies in mice have shown (*16*–*19*). However, NASH liver derived organoids exhibit a severe failure to dedifferentiate from the hepatic state back to the biliary state. Transcriptome analysis of all of our 6 NASH organoids show multiple consistently and strongly downregulated cell cycle pathways, a remarkable finding given their otherwise very heterogeneous transcriptomes, but entirely consistent with our functional data. Our data on NASH organoids are consistent with the well documented regenerative defect of NASH livers in both mice and humans, a problem of high medical relevance. For example, when tumors must be excised from NASH liver, the amount of liver tissue that can be removed is often significantly limited by the reduced capacity of a NASH liver to regrow.

iPSC – derived hepatic organoids have to our knowledge not been shown to have the capacity to repeatedly cycle between the biliary and hepatic states and further work is needed to explain this.

Regarding free fatty acid induced lipid droplet formation in NASH organoid flat cell outgrowth, we see a heterogeneity among NASH patients. While our NASH organoid transcriptomes do not show strong changes in lipid metabolism, this could be explained by the fact that the organoids we used for RNAseq have not been subjected to free fatty acid treatment, something that seems worth doing in the future. In addition the organoids used for transcriptome analysis were mostly 3D organoids with some flat cells. Indeed, despite our impression from visual inspection, we cannot rule out that the monolayers we are analyzing by microscopy react somewhat differently to oleic acid than the 3 D organoids in the same well. Thus, further study could include careful quantitation of 2D vs 3D areas in the same well by confocal microscopy and also total triglyceride measurement per well by chemical assay. To get an even more complete picture of potential differences between 2D and 3D areas, one could analyze the transcriptomes from both 2D and 3 D areas separately and at least some differences may be expected, such as in cell cycle control.

Two remarkable heterogeneities of the transcriptomes of NASH organoids are seen: First, NASH patients 1 and 5 show as their strongest upregulated pathways retinol metabolism, xenobiotic p450 related pathways and steroid hormone and porphyrine related pathways - indeed more than 10 fold. While patient 2 still shows a subset of these strongly upregulated pathways, patients 3, 4 and 6 show very little, if any, such upregulation. The reasons for this astounding difference are unclear, although we could imagine that some high cytochrome p450 pathway patients may have been subjected to drugs that elicit a strong cytochrome p450 response. However, it is remarkable that the p450 system is induced in patient organoids after 5-6 passages in organoid culture without the presence of any xenobiotics in the medium. Cleary more work is needed to interpret these data. The second heterogeneity consists in very strong upregulation of multiple proinflammatory pathways in patients 1,2,3 and 4 but only weak upregulation in patients 5 and 6. Human NASH livers are indeed characterized by high expression of pro inflammatory pathways, but the degree of heterogeneity among our NASH organoids is surprising given that all these organoids stem from end stage NASH patients with presumably high inflammation.

NASH patient liver derived organoids react in a remarkably patient specific way to pro regenerative / anti fibrotic drugs currently in clinical trials, such as obeticholic acid. This observation adds to the body of evidence that personalized treatment of NASH patients will be important in the future (*23*, *27*, *28*). Indeed, the study of NASH liver derived organoids may be helpful in this regard. In addition, HTP screens using a variety of different NASH organoids representing the different NASH subgroups may be inherently more powerful than HTP screens that are currently done with normal iPSC-Heps or iPSC-organoids. Patient specific HTP screens may also uncover novel classes of drugs that can alleviate steatosis, inflammation, fibrosis and cirrhosis in specific NASH subgroups.

## Materials and Methods

### Organoid Isolation

Liver pieces from both living and deceased normal donors and from end-stage NASH explanted liver were received from the University of Pennsylvania Transplant Center. Isolation was with modifications done according to (*12*, *13*). Liver pieces were stored at 4 degrees C in organoid basal medium (Advanced DMEM/F12, 1x Pen/Strep, 1x glutamax, 1x Hepes) for a maximum of 48 hours. Tissue samples were minced with a sterile razor blade in a sterile petri dish and then placed in digestion solution (EBSS, Collagenase D 2.5mg/mL, DNaseI 0.1mg/mL) and rocked at 37 degrees C for 30 to 90 minutes, depending on the sample. Patient explants needed generally about 60-90 min of collogenase digestion and donor biopsies usually 30 min. The tissue and cell solution was then washed with organoid wash medium (DMEM high glucose, 1x glutamax, 1x pyruvate, 1x pen/strep and 1%FBS) followed by centrifugation for 5 minues at 1300rpm.

### Organoid Spliting

To passage the organoid cultures, the basement matrix was removed by gently scraping the well to dislodge matrix droplet and incubating the drop in 500μL of basal medium on ice for 5 minutes. The cells were then pipetted up and down 20 times with a p1000 and a p10 tip attached to the end. Up to 3 50uL drops form a well of a 24 well plate can be pooled into the same tube. Additional basal medium was then added at a rate of 2mL per drop of matrix to further dissolve the matrix and the entire volume was mixed well by pipetting. After centrifugation for 5 minutes at 1300 rpm, the basal medium was carefully aspirated without disturbing the pellet. The pellet was resuspended in the desired volume of matrix (50μL drops for a 24 well plate, 5-15 μL drops for a 48 well plate) and seeded onto suspension culture plates. Drops were incubated 10-20 minutes at 37°C followed by overlay with prewarmed expansion medium according to (*12*, *13*) (43 mL Basal media supplemented with 1x B-27 supplement without vitamin A, NAC (1mM), Nicotinamide (10mM), 5mL RSPO-harvest medium, 1x N2 supplement, FGF-10 (100ng/mL), HGF (25ng/mL), human EGF (25ng/mL), leu15 Gastrin (10nM), Forskolin (10μM), and A83-01 (5μM). When seeding organoids, the expansion medium was supplemented with ROCK inhibitor Y27632 (10uM). For extended culture of ductal organoids, we omitted ROCK inhibitor in the medium.

### Hepatic Differentiation

Before differentiating human organoids, the expansion medium was supplemented with BMP-7 (100ng/mL) for approximately 5 days. Ductal stem cell organoids were then differentiated to hepatocyte like cells by the addition of a defined differentiation medium according to (*12*, *13*) which was composed of advanced Organoid Basal Medium (Advanced DMEM/F12, 10mM Hepes, 1x Glutamax, 1% pen/strep) supplemented with B-27 supplement (1x, with vitamin A), N2 supplement (1x), N-acetylcysteine (1mM), FGF-19 (100ng/mL), human EGF (50ng/mL), HGF (25ng/mL), leu15 gastrin (10nM), BMP-7 (25ng/mL), DAPT (10μM), dexamethasone (3μM), and A83-01 (500nM).

### Dedifferentiation of Hepatic Organoids

After 13 days in differentiation medium, organoid cultures were dedifferentiated. Each well was harvested in 500μL of cold basal medium in a sterile Eppendorf tube, pipeted up and down 30 times, and spun on a tabletop centrifuge at 3200rpm for 5 minutes. The basal medium was then removed and the cell pellet was resuspended in 50μL fresh, cold basal medium. 30μL of cell suspension was then divided into 3 sterile Eppendorf tubes (10μL each). 50μL of cold basal medium was added to each tube before spinning in a tabletop centrifuge at 3200rpm for 5 minutes. Basal medium was removed completely from each Eppendorf tube and the pellet was resuspended in 20uL of BME-2 matrix and distributed in 10μL drops on a 48 well suspension culture plate. Drops were solidified after a 15 minute incubation at 37 degrees C and overlaid with Expansion medium supplemented with ROCK inhibitor. The organoids were cultured in expansion medium for 12 days.

### Redifferentiation of Organoids

Dedifferentiated organoids were split into fresh BME-2 matrix and overlaid with expansion medium supplemented with ROCK inhibitor and BMP-7. When organoids had expanded to occupy most of the BME-2 matrix (approximately 5 days), the medium was exchanged for hepatic differentiation medium. Organoids were cultured in differentiation medium for 13 days before imaging and harvesting RNA.

### RNA Isolation from Hepatic Organoids

RNA was harvested from 3 individual wells on a 24 well plate for each of 10 organoid populations. For all lines, 3 wells were seeded in 50μL of BME-2 matrix and differentiated after approximately 5 days of expansion. Each well was harvested individually after 13 days of differentiation by removing the medium and adding 500uL of TRIZOL to the well. The entire matrix drop and any flat cells was resuspended in the TRIZOL by gentle scraping of the bottom and then transferred to an RNase free 1.5mL eppendorf tube. Sampels were flash frozen on dry ice and stored at −80 degrees Celcius.

Samples were then thawed, treated with 100uL of chloroform, inverted several times and incubated at room temperature for 1-2 minutes before centrifugation at 13,000 rpm for 20 minutes. The aqueous upper phase was then transferred to a new RNase free Eppendorf tube for clean up using Qiagen’s RNeasy Mini Kit with the on - column DNase digestion protocol. The concentration of each sample was measured using the Take3 trio microspot plate and Biotek plate reader. Integrity of the RNA was assessed by running 4uL of eluted product on an RNase free 0.8% agarose gel.

### Preparation of RNA Libraries for RNAseq

50ng of purified RNA for each sample was transferred to a clean 96 well PCR plate for mRNA library preparation. Each cDNA library was synthesized using the NEBNext Ultra II Directional RNA Library Prep Kit (catalog# e7765) with the NEBNext Poly(A) mRNA Magnetic Isolation Module (catalog# e7490), and NEBNext Multiplex Oligos for Illumina (catalog# e6609) according to manufacturer’s instructions.

11 samples were run on a high sensitivity DNA chip through a bioanlyzer to determine representative average length of products in each library. The molarity of each library was then analyzed using the KAPA Biosystems ROX low qPCR complete kit (ROCHE cat# 07960336001). All libraries were then transferred to the UPenn Next Generation Sequencing Core for analysis on an Illumina HiSeq 4000 analyzer.

### LDL Uptake Analysis

4 wells of each organoid cell line were seeded in BME-2 matrix on suspension 48 well plates and overlaid with expansion medium supplemented with ROCK inhibitor and BMP-7. After organoids occupied most of the matrix droplet (approximately 5 days), the medium was changed to differentiation medium. On day 13 of differentiation, LipidTOX live cell lipid stain was added 1:1000 to each well and incubated overnight. Representative images were taken using NIS elements software (Nikon) on day 14 of differentiation before adding PHRODO green LDL stain 1:500 and incubating overnight. After the cells have been double labeled, 9 representative image series were taken of the flat cell monolayers extending outside the BME drop. Each image series consisted of the LDL uptake, Lipid accumulation, and in some cases phase contrast filters for the same field of vision. LDL uptake was quantified using Nikon NIS elements object count software with universal diameter settings of 1-50um and universal circularity settings of 0-1. Total cell area was measured using the NIS elements area tool on strongly over exposed lipidTOX images that clearly reveal all cell areas, even those with very little lipid accumulation.

### Free Fatty Acid Uptake Analysis

On 48 well suspension plates, 8 wells of each patient or donor organoid line were seeded in BME matrix and overlaid with expansion medium supplemented with ROCK inhibitor and BMP-7. Organoids were grown for 6-11 days (depending on the donor or NASH patient) before exchanging the medium for hepatic differentiation medium. On day 12 of differentiation, differentiation medium was supplemented with either 15mg/mL fatty acid free BSA and 2mM Oleic Acid or 15mg/mL BSA only. A stock of BSA was prepped by dissolving BSA in PBS without calcium or magnesium to a final concentration of 150mg/mL. 82mM oleic acid dissolved in methanol was supplied by the Rader lab (D.M.C). The BSA/PBS solution was added directly to a large aliquot of differentiation medium to a final concentration of 15mg/mL and divided in half. To one aliquot of BSA-differentiation medium, oleic acid was added to a final concentration of 2mM.

For each organoid line, 2 wells were treated with 200uL of BSA-differentiation medium and 2 wells were treated with 200uL of BSA-Oleic acid-differentiation medium for 24 hours. After a 24 hour incubation, each well was stained with 1:1000 LipidTOX, 1:1000 cellEvent caspase 5/7 detection dye, and 2uM Hoechst dye. All dyes were prepared together as a 3x concentrate in differentiation medium of which 100uL was added to each experimental well. All wells were incubated for an additional 4 hours.

Immediately before microscopy analysis, the differentiation medium was changed to remove any unintegrated dyes and BSA. 10 image series were taken for each organoid population in both +oleic acid and −oleic groups. 1 image series consisted of lipidtox detection (red), cellEvent detection (green), and Hoechst dye (blue) for the same visual field. Sum intensity of LipidTOX images was measured using NIS elements software. Cell number was determined from Hoechst dye images using NIS elements software.

---

**Supplemental Figure 1.**
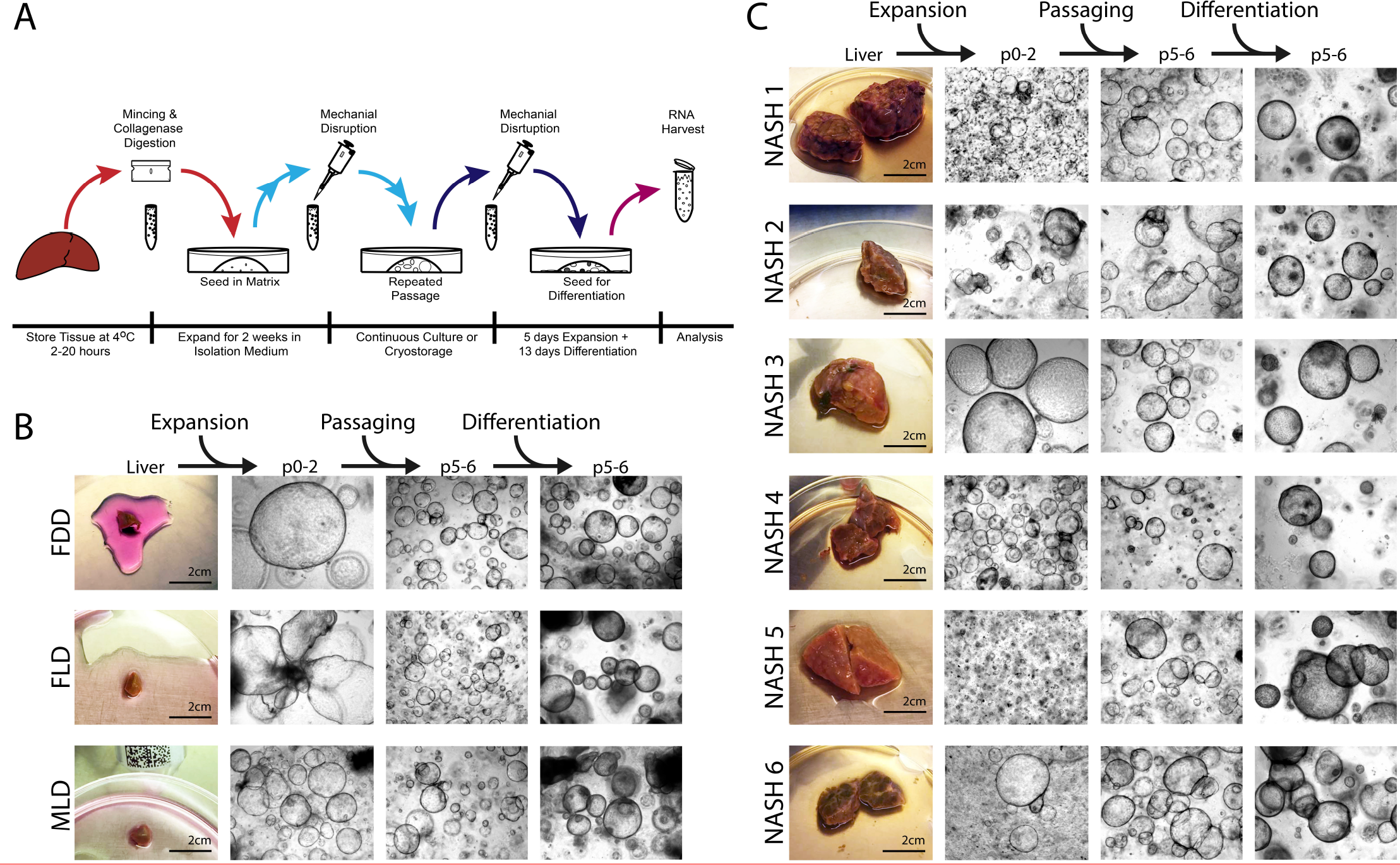
Ductal organoids isolated from human explant liver grow more slowly than those from normal liver biopsies. **A.** Schematic of isolation of bipotent ductal organoids from human liver. Viable bipotent ductal organoids were isolated from all samples by mincing them with razor blades, followed by collagenase digestion and embedding in BME matrix under expansion medium. Organoids were allowed to develop for about 2 weeks and subsequent passaging was performed by mechanical disruption and seeding into BME matrix with expansion medium. All organoids were further passaged in expansion medium 5-6 times. Average times between splits was about 3-5 days. 4 days after the last split organoids were differentiated to the hepatic state by switching to differentiation medium. At day 13 of differentiation organoids were harvested for isolation of total RNA using Trizol reagent. **B.** Bipotent Ductal Organoids were isolated from 50-100mg of normal donor liver. **C.** Bipotent Ductal Organoids were isolated from either 300-500mg or only 100mg of NASH patient explants. The rest of the explant tissue was frozen down for later use in genomics and transcriptomics. Viable bipotent ductal organoids were isolated from all biopsies and explants shown in B and C.

**Supplemental Figure 2.**
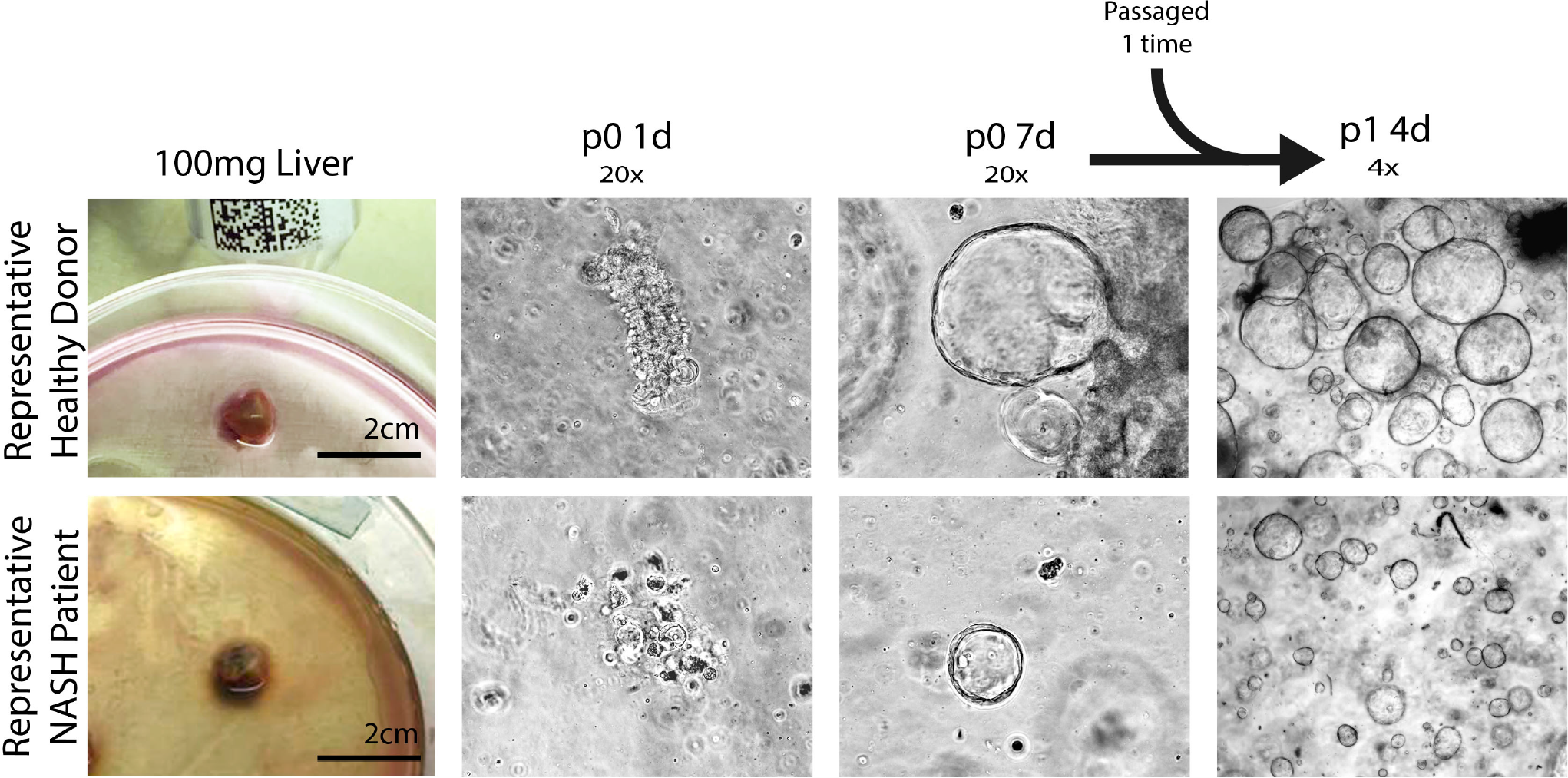
Organoids derived from most human NASH explant liver samples exhbit a significant developmental delay compared to organoids derived from normal human liver. Bipotent ductal organoids were isolated in parallel from 100mg human donor liver and 100mg of the corresponding NASH liver explant. Representative end stage NASH explant organoids exhibit a significant developmental delay compared to healthy donor organoids. NASH liver organoids demonstrate slower growth and smaller organoid size at p0 after 7d of culture and p1 after 4 days of culture.

**Supplemental Figure 3.**
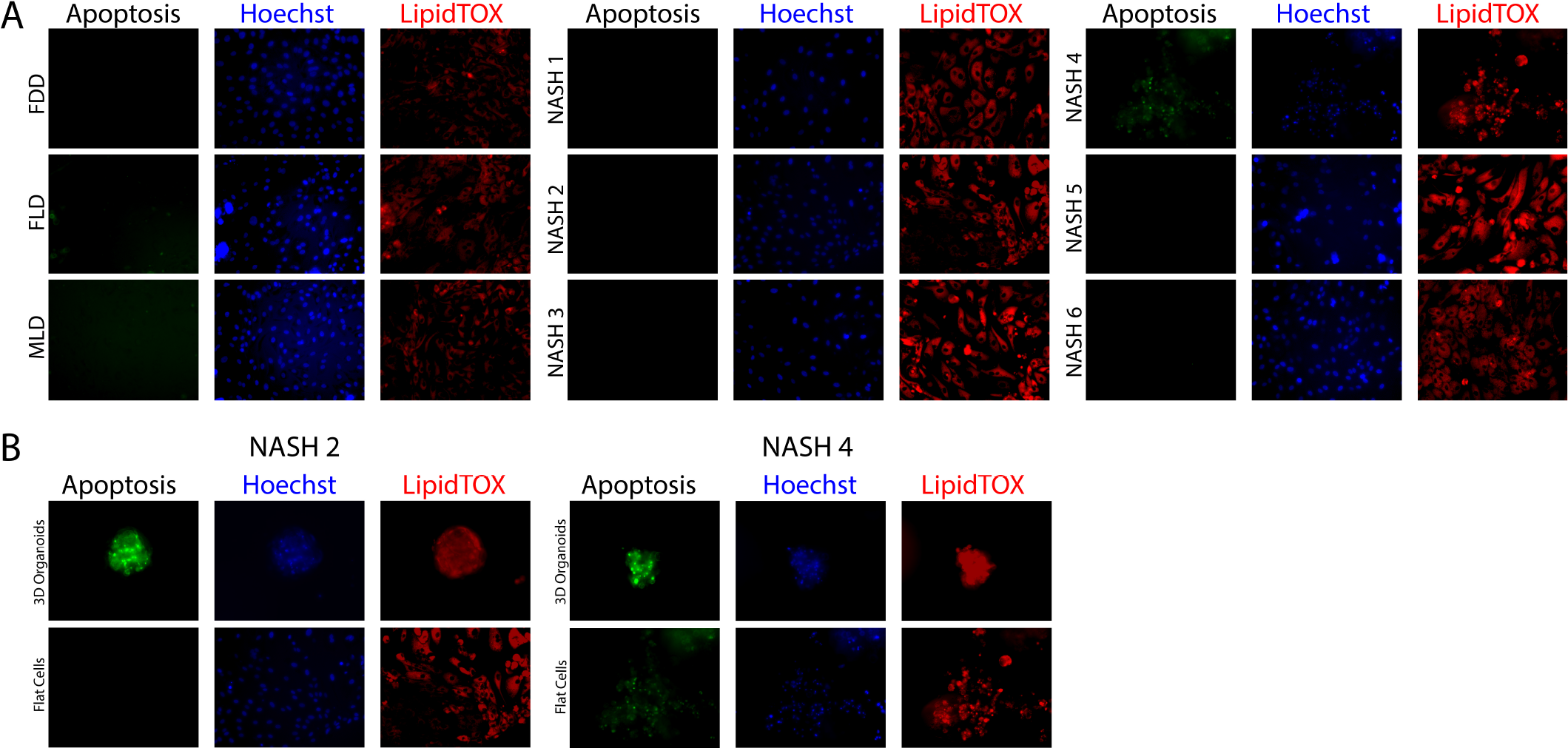
Hepatically differentiated organoids isolated from human NASH liver explant grown as monolayers exhibit low levels of apoptosis. **A.** On day 13 of differentiation organoids from normal and NASH donors were triple labeled with CellEvent (a caspase 3/7 indicator), Hoechst and LipidTOX. 2mM of oleic acid was added to all organoid populations. After overnight incubation flat cells were imaged. Representative images of monolayers (from a total of 10 images) for each of the three donors and each of the 6 NASH patients are shown. **B.** 3D organoids from the same culture well that also contained the monolayers shown in A (for patients NASH2 and NASH4 only) were imaged.

## Acknowledgments

We thank Joseph Kutch, Shilpa Rao and Jonathan Schug from the Penn SOM Next Generation Sequencing Core for advice on preparation, quality control and data analysis of RNAseq libraries and Jinlong Feng for facilitating transport and cold storage of the liver biopsies and explants.

## Author contributions

S.M. and B.B. performed most experiments, prepared the figures and edited the manuscript, D.A. and D.M.C contributed to lipid staining, free fatty acid induced lipid accumulation and other lipid related procedures and edited the manuscript, D.J.R. edited the manuscript, K.O. and A.S. procured the liver biopsies and liver explants based on their expertise as transplant surgeons, and edited the manuscript, T.D.R. established the research strategy, designed most experiments, performed all derivations of organoids from liver biospies and explants, and wrote and edited the manuscript.

